# Disrupting the CD177:proteinase 3 membrane complex reduces anti-PR3 antibody-induced neutrophil activation

**DOI:** 10.1101/2021.05.17.444335

**Authors:** Stephen F. Marino, Uwe Jerke, Susanne Rolle, Oliver Daumke, Ralph Kettritz

## Abstract

CD177 is a neutrophil-specific receptor presenting proteinase 3 (PR3) autoantigen on the neutrophil surface. CD177 expression is restricted to a neutrophil subset resulting in CD177^pos^/mPR3^high^ and CD177^neg^/mPR3^low^ populations. The size of the CD177^pos^/mPR3^high^ subset has implications for anti-neutrophil cytoplasmic autoantibody (ANCA)-associated autoimmune vasculitis (AAV) where patients harbor PR3-specific ANCA that activate neutrophils. We generated high affinity anti-CD177 monoclonal antibodies, some of which interfered with PR3 binding to CD177 (PR3 “blockers”) as determined by surface plasmon resonance spectroscopy, and used them to test the effect of competing PR3 from the surface of CD177^pos^ neutrophils. Because intact anti-CD177 antibodies also caused neutrophil activation, we prepared non-activating Fab fragments of a PR3 blocker and non-blocker that bound specifically to CD177^pos^ neutrophils by flow cytometry. We observed that Fab blocker clone 40, but not non-blocker clone 80, dose-dependently reduced anti-PR3 antibody binding to CD177^pos^ neutrophils. Importantly, preincubation with clone 40 significantly reduced respiratory burst in primed neutrophils challenged either with monoclonal antibodies to PR3 or PR3-ANCA IgG from AAV patients. After separating the two CD177/mPR3 neutrophil subsets from individual donors by magnetic sorting, we found that PR3-ANCA provoked significantly more superoxide production in CD177^pos^/mPR3^high^ than in CD177^neg^/mPR3^low^ neutrophils, and that anti-CD177 Fab clone 40 reduced the superoxide production of CD177^pos^ cells to the level of the CD177^neg^ cells. Our data demonstrate the importance of the CD177:PR3 membrane complex in maintaining a high ANCA epitope density and thereby underscore the contribution of CD177 to the severity of PR3-ANCA diseases.

---

As the most abundant leukocytes, neutrophil granulocytes represent one of the first lines of defense against infectious agents and are therefore a pillar of the innate immune system. Among their most potent defense mechanisms are the respiratory burst to generate reactive oxygen species (ROS) and degranulation, whereby stores of cytotoxic species housed in several types of intracellular, membrane bound compartments called granules, are moved to the cell surface and released into the extracellular environment as a response to pathogen detection (1). This toxic cocktail is designed to kill foreign cells in the vicinity of the neutrophil. Given that healthy cells are also negatively affected, degranulation is a highly regulated process (though not yet fully understood) (2). The serine protease proteinase 3 (PR3) is found in large abundance in human neutrophils (3). It is a major component of neutrophil azurophilic granules but is interestingly also detectable on the outer surface of the neutrophil plasma membrane. In most individuals, two distinct neutrophil populations can be identified based on the amount of membrane bound PR3 (mPR3) they harbor – one with low amounts of mPR3 (mPR3^low^) and another with orders of magnitude more detectable mPR3 (mPR3^high^) (4). The mPR3^high^ population is further distinguished by the presence of a selectively expressed membrane receptor called CD177 (5,6). CD177 is a GPI-anchored protein exclusively expressed in a subset of neutrophils and forms a high affinity complex with PR3. It thus accounts for the increased mPR3 levels that are detectable on the mPR3^high^ subset (7). The proportion of CD177^pos^/mPR3^high^ vs CD177^neg^/mPR3^low^ neutrophils in a given individual is genetically determined and remains constant throughout life (8-10). Although the function of CD177 is still unclear, several studies have identified a correlation between a large CD177^pos^/mPR3^high^ neutrophil population and the occurrence and progression of a group of incurable autoimmune diseases called ANCA (<u>a</u>nti-<u>n</u>eutrophil <u>c</u>ytoplasmic <u>a</u>ntibody) vasculitides (8,11-14). In these disorders, autoantibodies directed against PR3 stimulate respiratory burst and degranulation. The resulting release of ROS and cytotoxic enzymes and peptides – circumventing the normally strictly controlled degranulation process - causes considerable systemic damage to healthy tissue and is the hallmark of these conditions. It has been shown that, although all neutrophils are activated upon exposure to PR3-ANCA, CD177^pos^/mPR3^high^ neutrophils react more strongly to autoantibody binding, as measured by degranulation, generation of superoxide, an initial product of the respiratory burst (referred to as ‘oxidative burst’), and increased phosphorylation of Akt kinase (15). AAV patients with large CD177^pos^/mPR3^high^ populations are more prone to relapse and show poorer clinical outcomes than those with smaller CD177^pos^/mPR3^high^ populations (11-13).

The mechanism by which PR3-ANCA cause neutrophil activation is not known. Since all neutrophils display mPR3 and are affected by PR3-ANCA, the presence of PR3 seems critical for the process. In the case of CD177^pos^/mPR3^high^ neutrophils, which are more strongly affected by the binding of PR3-ANCA, the questions arise whether and how CD177 itself may contribute to ANCA-stimulated degranulation. Although CD177 does not cross the plasma membrane, it could interact with other species that do and in this way enhance the sensitivity of CD177^pos^/mPR3^high^ neutrophils to the effects of PR3-ANCA.

We sought to directly test the contribution of CD177 to PR3-ANCA-stimulated neutrophil activation. To this end, we generated a series of anti-CD177 antibodies, some of which bound to the CD177:PR3 complex and some of which blocked the binding of PR3. We used Fab fragments derived from the latter to selectively disrupt CD177:PR3 complexes on CD177^pos^/mPR3^high^ neutrophils. We then tested the effect of this treatment on PR3-ANCA induced respiratory burst using both mixed and sorted neutrophil populations. We show that removing CD177 bound PR3 reduces the sensitivity of mixed neutrophil pools to PR3-ANCA treatment. When we tested separated CD177^neg^/mPR3^low^ and CD177^pos^/mPR3^high^ populations, we found that while Fab treatment had no effect on PR3-ANCA-induced respiratory burst of CD177^neg^/mPR3^low^ neutrophils, the anti-CD177 Fabs reduced the response of the CD177^pos^/mPR3^high^ population to that of the CD177^neg^/mPR3^low^ population. Thus, the excess mPR3 on CD177^pos^/mPR3^high^ neutrophils appears to account for their enhanced sensitivity to PR3-ANCA. The presence of CD177 enables a higher density of PR3 epitopes that result in a stronger activation effect in response to autoantibody binding than seen in CD177^neg^ neutrophils.

## Results

### Screening of anti-CD177 monoclonal antibodies identifies binders that block the CD177:PR3 interaction

We used recombinant CD177 (7) for the generation of mouse monoclonal antibodies against CD177. 10 of the resulting hybridoma products were assessed by surface plasmon resonance (SPR) spectroscopy to determine their binding affinities for CD177. All 10 IgGs bound CD177 with high affinity, ranging from 0.14 to 11.9 × 10^−9^ M (Table 1). We attempted to determine the ability of each IgG to block the binding of PR3 to CD177 by competition ELISA using an anti-PR3 antibody but were unable to optimize the assay to produce unambiguous results. We therefore devised a direct assay using SPR as depicted in Fig. 1. We immobilized each anti-CD177 IgG to the sensor chip and subjected them to two back-to-back ligand flows: the first ligand sample contained only CD177, which was allowed to flow over the immobilized IgGs until the resonance units (RUs) indicated near saturation of all binding sites (Figure 1A, B); the second ligand sample containing CD177:PR3 complexes was then injected and the resulting effect on the RUs observed. Two possible RU responses could be expected. If the IgG in question does not interfere with the PR3 interaction with CD177 (i.e. the IgG was a *non-blocker*), then CD177:PR3 complexes could be exchanged for CD177 in the IgG binding sites, leading to a second pronounced increase in the RUs as the extra mass from PR3 was added to each binding interaction (Fig. 1B). Conversely, if the IgG does prevent interaction of PR3 with CD177 (i.e. the IgG is a *blocker*) then no further RU increase would be possible, but rather a decrease in RUs during flow of the complex, since bound CD177 will be lost from the immobilized IgG and there is insufficient free CD177 in the complex mixture to take its place. To minimize the concentration of free CD177 in the complex sample, the concentration used was 10-fold higher than the PR3 binding affinity, as measured also by SPR ((7), Fig. 1B). Using this assay, we unambiguously identified three PR3 blockers among the tested IgGs (Fig. 1C, clones 7, 40 and 72), while the remainder were non-blockers (Table 1).

**Table 1:** Summary of the binding affinities and complex blocking abilities of the anti-CD177 monoclonal antibodies.

| Blocker? |  |  |  |
| --- | --- | --- | --- |
| Antibody | Affinity | ELISA | SPR |
| 7-3-1 | $8.0 \times 10^{-9}\text{M}$ | yes | yes |
| 39-5-2 | $0.14 \times 10^{-9}\text{M}$ | no | no |
| 40-6-4 | $3.1 \times 10^{-9}\text{M}$ | yes | yes |
| 49-3-1 | $11.9 \times 10^{-9}\text{M}$ | ~yes | no |
| 72-3-1 | $9.0 \times 10^{-9}\text{M}$ | yes | yes |
| 73-8-1 | $0.92 \times 10^{-9}\text{M}$ | ~yes | no |
| 80-11-2 | $2.3 \times 10^{-9}\text{M}$ | ?? | no |
| 82-2-4 | $0.76 \times 10^{-9}\text{M}$ | ~yes | no |
| 90-12-1 | $1.3 \times 10^{-9}\text{M}$ | ~yes | no |
| 92-3-4 | $11.8 \times 10^{-9}\text{M}$ | ~yes | no |

**Figure 1:**
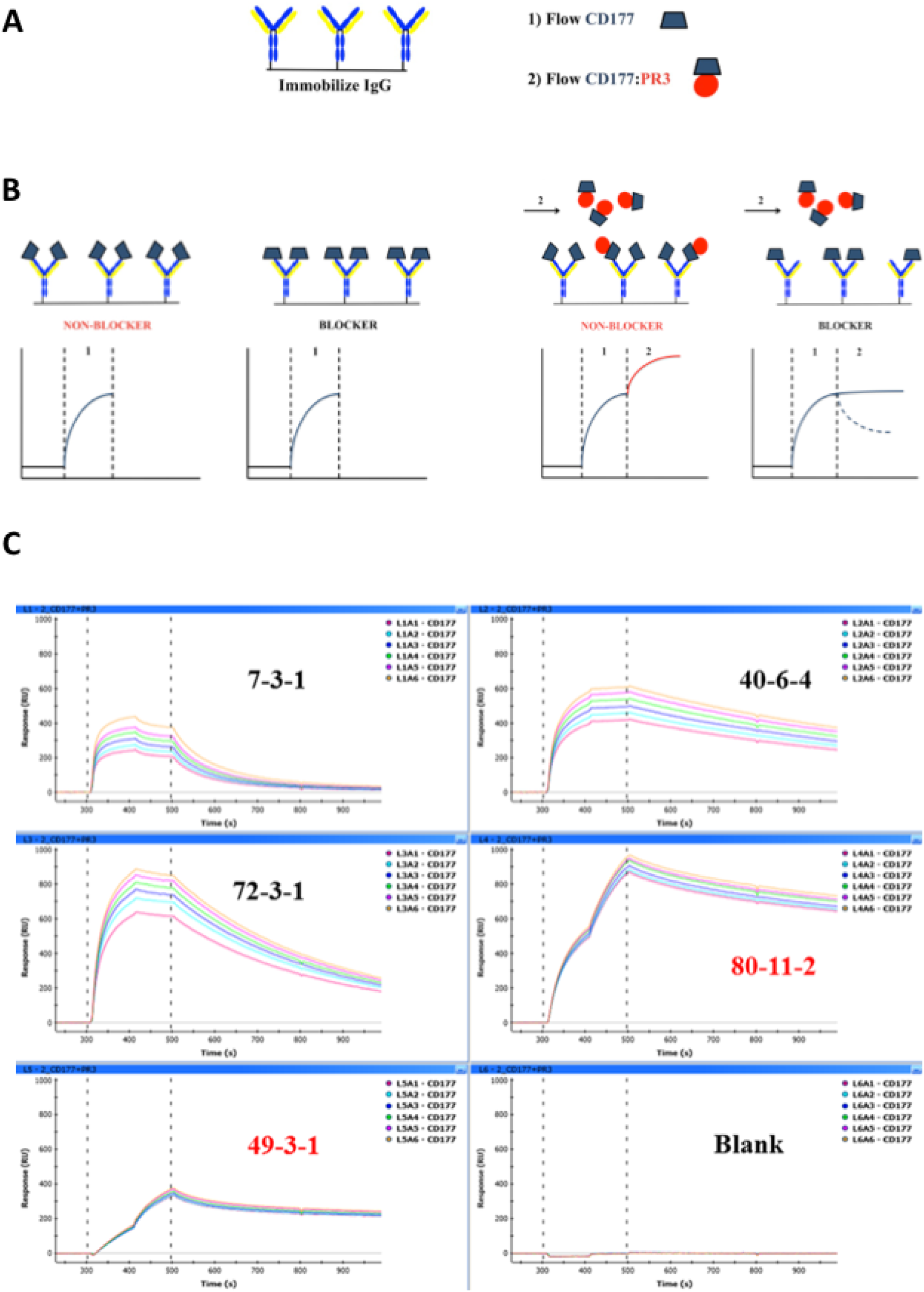
(A) IgG was immobilized on the SPR sensor chip and subjected to two consecutive ligand flows – first with CD177 alone and second with the CD177:PR3 complex. (B) Expected sensorgrams for each ligand flow for both blocker and non-blocker IgG; left panel, ligand flow 1; right panel, ligand flow 2. (C) SPR sensorgrams showing blocker (labeled in black) and non-blocker (labeled in red) IgG; Ligand flows 1 and 2 occurred between the horizontal dotted lines.

### PR3 blocker and non-blocker IgGs and their corresponding Fab fragments bind to CD177^pos^ neutrophils, but only IgGs are activating

We next verified that our anti-CD177 monoclonal antibodies could bind to intact human CD177^pos^ neutrophils. Total neutrophil pools isolated from freshly donated blood samples were incubated with purified anti-CD177 IgG and subjected to FACS analysis. Binding was confirmed in all cases (data not shown).

We also tested purified Fab fragments derived from each IgG by papain digestion and, again, confirmed binding of each to CD177^pos^ neutrophils by FACS staining (Fig. 2A).

**Figure 2:**
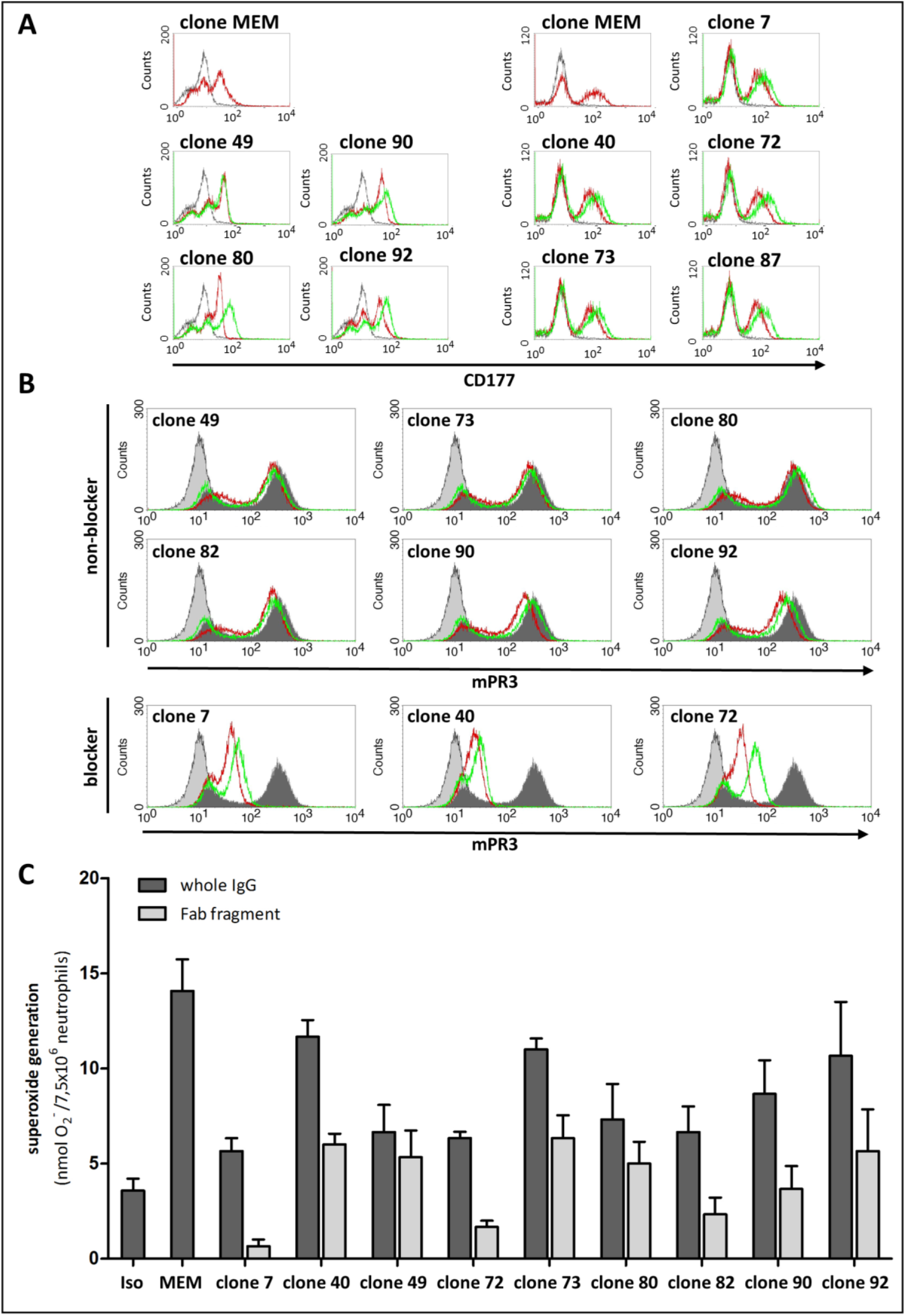
(A) Freshly isolated Neutrophils were incubated with 5 µg/ml anti-CD177 IgG (red line) and anti-CD177 Fab (green line) followed by incubation with a FITC conjugated goat anti-mouse IgG (Fab specific (Sigma, #F5262) secondary antibody. Isotype staining is shown in grey. Shown experiments were perfomed in two sets with two different human blood donors (B) Neutrophils were primed with TNFa (2ng/ml) and incubated with non-blocker or blocker anti-CD177 IgG or corresponding Fab (20µg/ml), following incubation with anti-PR3-Alexa488 IgG (2,5µg/ml). Histograms show isotype staining (light grey), mPR3 staining (dark grey), and the effect of anti-CD177 IgG (red line) and anti-CD177 Fab (green line) on the mPR3 staining accordingly. (C) Superoxide generation in neutrophils stimulated with anti-CD177 whole IgG (5 µg/ml) (dark gray bars) or corresponding Fab (5 µg/ml) (light grey bars). A non-CD177 binding isotype was used as negative control (Iso), a commercially available, activating monoclonal IgG against CD177 (clone MEM) was used as positive control (MEM). n=3

Next, neutrophils were preincubated with our anti-CD177 IgGs and/or Fab fragments to test if PR3 blockers (clones 7, 40, and 72) interfered with subsequent anti-PR3 IgG binding (Fig. 2B). In contrast to PR3 non-blockers, the PR3 blockers reduced the anti-PR3 staining signal in the CD177^pos^ neutrophil subset.

Multivalent PR3 binders - like PR3-ANCA – strongly activate neutrophils for respiratory burst and partial degranulation, but monovalent binders – like Fab fragments – do not (16). We therefore tested whether our CD177 binders also produced respiratory burst. We incubated isolated primed neutrophils with either anti-CD177 IgG or their corresponding Fabs and then determined to what extent they initiated oxidative burst by measuring the production of superoxide in the resulting aliquots. Though to differing extents, in all cases the multivalent anti-CD177 IgG provoked respiratory burst in mixed neutrophil pools (Fig. 2C). Their corresponding monovalent Fabs, however, showed no significant stimulatory effect, with most of the resulting superoxide concentrations comparable to the negative control value. These effects also showed no obvious correlation with either the blocking or non-blocking properties of the individual binders or their relative affinities to CD177. For all further experiments with neutrophils, we chose to proceed with the IgG blocker clone 40 (K_d_ = 3.1 × 10^−9^ M) and the non-blocker clone 80 (K_d_ = 2.3 × 10^−9^ M) since both binders have a CD177 affinity similar to that of PR3 (K_d_ = 4.1 × 10^−9^ M).

### A PR3 blocking Fab reduces anti-PR3 IgG stimulated oxidative burst in mixed (CD177^pos^/mPR3^high^ / CD177^neg^/mPR3^low^) neutrophil populations

We next tested whether preincubation of mixed population neutrophils with blocker Fab clone 40 had any effect on the stimulation of superoxide production by anti-PR3 IgG. We first incubated unsorted neutrophils with a saturating amount (20 μg/ml) of either blocker clone 40 or non-blocker clone 80 and, after washing, activated the neutrophils by addition of a monoclonal anti-PR3 antibody. The subsequent superoxide measurements showed that the anti-PR3 monoclonal IgG strongly stimulated the neutrophil pool in all cases, as opposed to treatment with an isotype IgG that provoked only a background response (Fig. 3A). While the degree of superoxide generation in the presence of non-blocker clone 80 was not significantly different than that measured in the absence of Fab or the presence of a non-CD177 binding control Fab, the pool containing blocker clone 40 showed a clearly weaker stimulation and correspondingly less superoxide production, implying that the PR3 blocking effect of clone 40 protects CD177^pos^ neutrophils from activation by anti-PR3 IgG. FACS staining performed in parallel showed that pretreatment with blocker clone 40 significantly reduced anti-PR3 antibody binding to the CD177^pos^ neutrophil subset, whereas non-blocker and control Fab showed no reduction in mPR3 expression (Fig. 3A, B).

**Figure 3:**
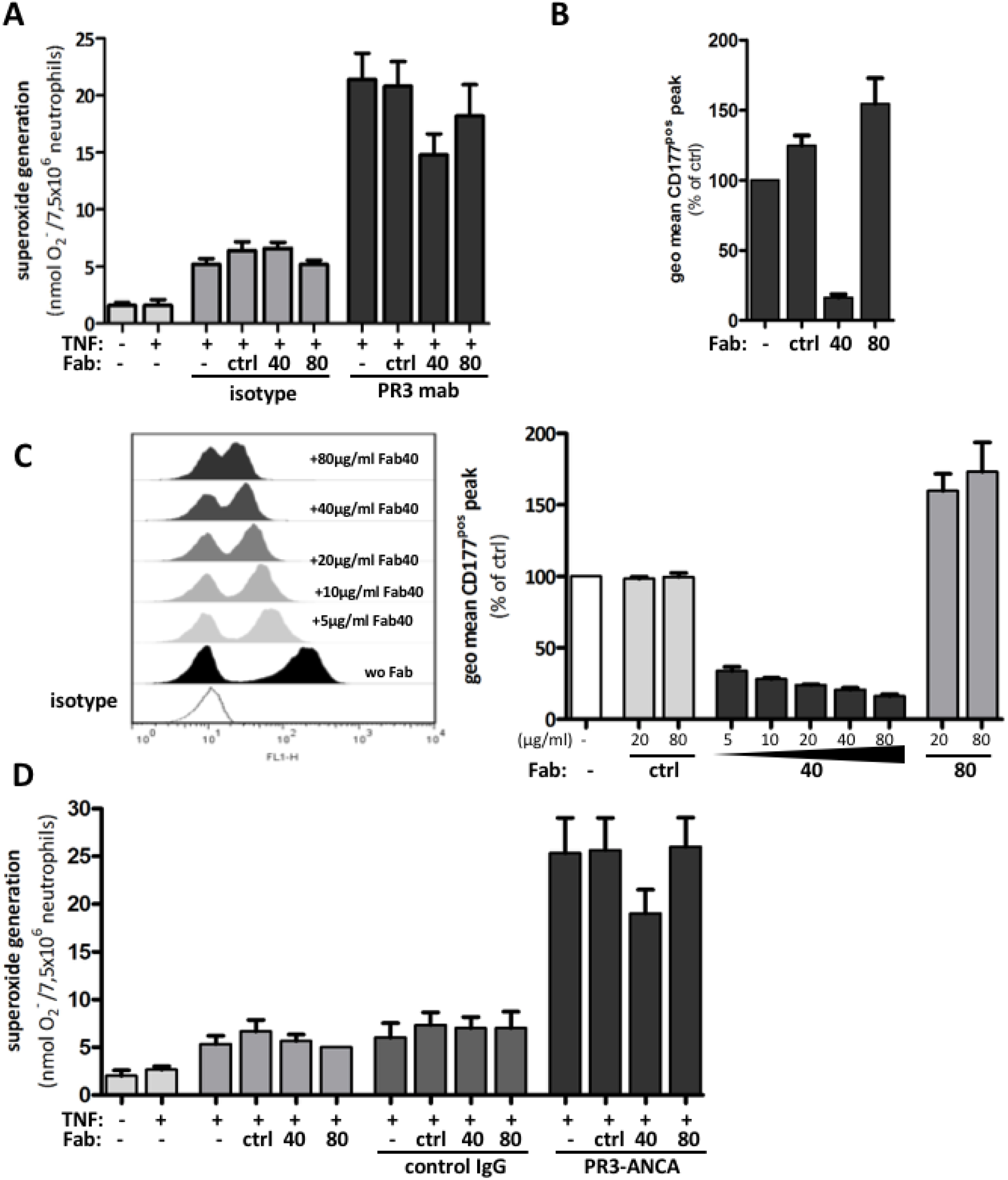
(A) Reduced superoxide production in unsorted neutrophils after incubation with blocking anti-CD177 Fab. Primed neutrophils were untreated or preincubated with 20 µg/ml control Fab (ctrl), blocker Fab (clone 40), or non-blocker Fab (clone 80), followed by 5 µg/ml stimulating monoclonal anti-PR3 IgG (dark grey bars), Isotype control (mid grey bars), or without any further stimulation (light grey bars). (left panel, n=5). In parallel mPR3 staining of the CD177^pos^ neutrophil subset was assayed after incubation with Fabs as described above. (right panel, n=5) (B) Dose–dependent reduction in mPR3 staining on neutrophils after incubation with blocking anti-CD177 Fab (clone 40). A representative set of histograms (left panel) and the corresponding geometric mean of mPR3 staining of the CD177^pos^ peak (right panel) from all experiments are shown. Untreated neutrophils were used as control. Non-CD177 binding control Fab (light grey bars), Fab clone 40 (dark grey bars), non-blocking Fab clone 80 (mid grey bars). (n=3). (C) Blocker Fab clone 40 negatively influences superoxide generation in neutrophils after PR3-ANCA stimulation (75 µg/ml). (n=3) Legend as in (A). Comparison between multiple groups were done using ANOVA and Bonferroni’s post-hoc test, * indicates p < 0.05

To verify that the blocker clone 40 Fab was displacing CD177 bound PR3 from the CD177^pos^ neutrophils, we incubated unsorted neutrophils with increasing concentrations of clone 40 Fab and monitored changes in detectable mPR3 by FACS. In the absence of Fab, the peaks corresponding to the CD177^neg^/mPR3^low^ and CD177^pos^/mPR3^high^ populations were widely separated; upon addition of the clone 40 Fab, the separation between the peaks (stained for PR3) decreased dose-dependently until they merged, with very little difference distinguishable between them at the highest Fab concentration tested (Fig. 3C). This observation confirmed that the blocker Fab 40 not only prevented PR3 binding to CD177 in SPR but also on the surfaces of living neutrophils. We then repeated our activation experiment with PR3-ANCA obtained from AAV patient serum. As in the previous experiment with an anti-PR3 monoclonal IgG, only preincubation with blocker Fab 40 had a negative influence on the stimulation of superoxide production by PR3-ANCA (Fig. 3D).

### The effects on neutrophil activation demonstrated by Fab blocker clone 40 are restricted to the CD177^pos^/PR3^high^ neutrophil population

Preincubation of unsorted neutrophils with the PR3 blocking clone 40 Fab not only displaced PR3 from the CD177^pos^ population, but also effected a clear reduction in the amount of superoxide produced as a result of stimulation with either monoclonal anti-PR3 IgG or PR3-ANCA IgG isolated from AAV patient serum. In order to more precisely define the effect of blocker clone 40, we repeated the PR3-ANCA stimulation experiments with sorted neutrophils. We used magnetic cell sorting with our isolated neutrophils to produce pure CD177^neg^/mPR3^low^ and CD177^pos^/mPR3^high^ preparations (Fig. 4A) and tested them separately by preincubation with 20 μg/ml blocker clone 40 Fab before washing and addition of the stimulatory PR3-ANCA. As previously shown (15), both pure populations were activated by the addition of PR3-ANCA, with the CD177^pos^/mPR3^high^ population producing nearly twice as much superoxide as the CD177^neg^/mPR3^low^ population (Fig. 4B). Preincubation with non-blocker clone 80 showed no effect on superoxide production; the values for both populations in the presence of this Fab were identical to those either without added Fab or preincubated with a control Fab. Preincubation with blocker clone 40 showed no measurable effect on superoxide production by stimulated CD177^neg^/mPR3^low^ neutrophils but a substantial effect with CD177^pos^/mPR3^high^ neutrophils. The presence of clone 40 completely eliminated the difference in superoxide production between the two pure neutrophil populations.

**Figure 4:**
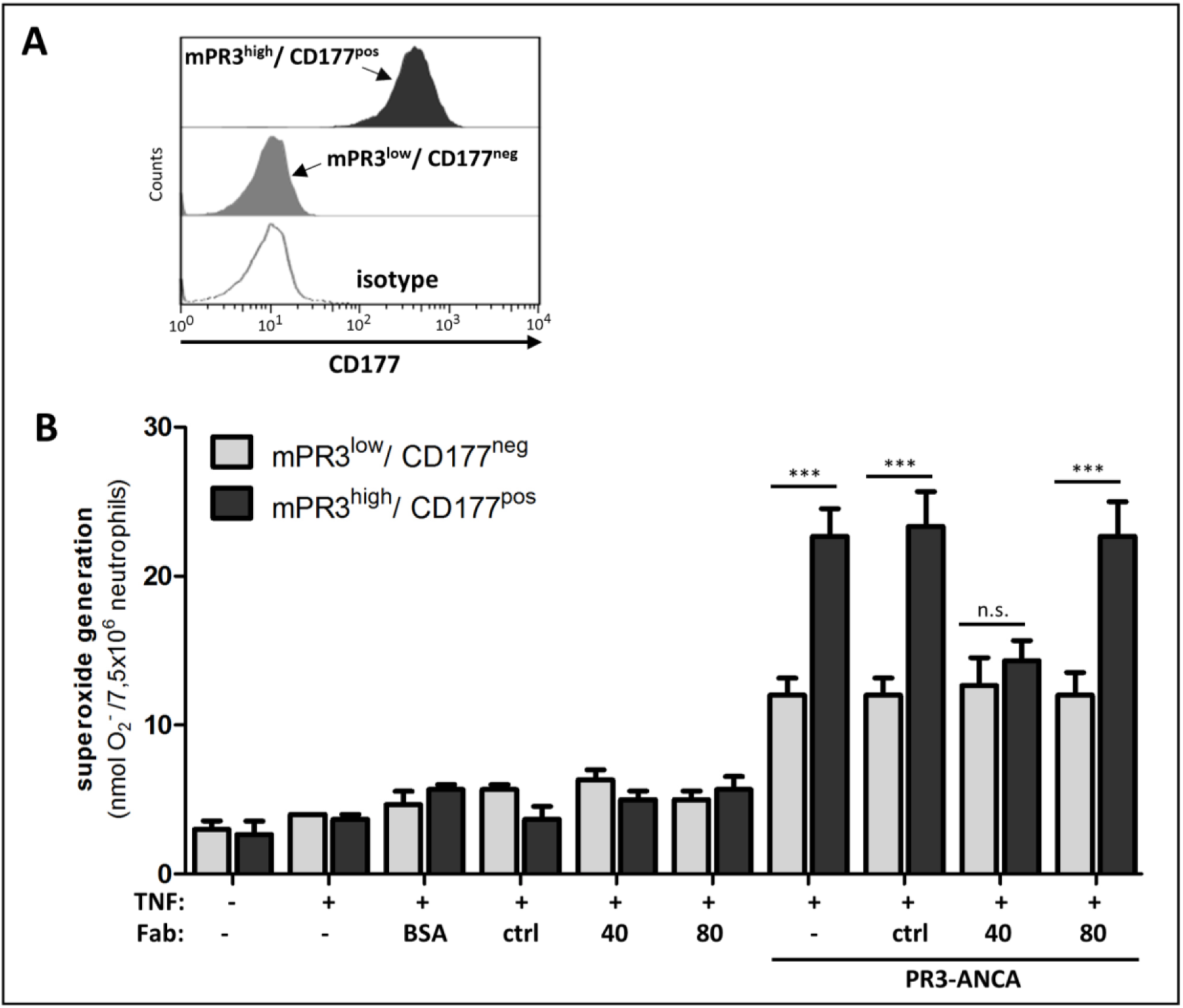
**(A)** Neutrophils from CD177/mPR3 bimodal donors were separated by magnetic cell sorting. Panel shows a representative separation after sorting into CD177^pos^/mPR3^high^ and CD177^neg^/mPR3^low^ subsets. **(B)** Blocking of neutrophil activation by anti-CD177 Fab clone 40 is restricted to the CD177^pos^/mPR3^high^ neutrophil subset. As described in Fig. 3A, sorted neutrophils (CD177^pos^/mPR3^high^, dark grey bars and CD177^neg^/mPR3^low^, light grey bars) were assayed for superoxide generation. Because a PR3-ANCA epitope could be blocked by the monoclonal PR3 antibody used for the sorting procedure, polyclonal human PR3-ANCA (75 µg/ml) were used for stimulation. (n=3) Comparison between multiple groups were done using ANOVA and Bonferroni’s post-hoc test, * indicates p < 0.05

## Discussion

PR3 is a member of a family of neutrophil serine proteases important in inflammation and is known to process extracellular matrix proteins (3), cell surface receptors (17,18), cytokines (19,20) and intracellular effectors including kinase inhibitors (21), cytoskeletal proteins (22), and annexin (23). As a major component of azurophilic granules, it is also released in abundance from neutrophils during degranulation. Although it is a soluble protein, it is detectable on the extracellular surface of all neutrophils and readily adheres to the membranes of non-myeloid cells as well (7). PR3 possesses a patch of hydrophobic residues on its surface that have been suggested to be responsible for its interaction with membranes (24), though definitive proof for this supposition has yet to be reported. To date, the only confirmed (non-substrate) interaction partner for PR3 is CD177, a GPI anchored protein with no clearly defined function. Due to its high affinity complex with PR3, CD177 is responsible for the abundant mPR3 occurring in neutrophils expressing it. As the target of autoantibodies, mPR3 is a central player in ANCA vasculitis that, in an as yet unknown manner, facilitates autoantibody induced respiratory burst and degranulation, a process normally requiring multiple, receptor initiated, signaling cascades (2). An outstanding question here is exactly how molecules that do not obviously function as receptor ligands and do not themselves cross the plasma membrane can nevertheless activate the intracellular signaling pathways necessary for the degranulation process. The fact that CD177^pos^ neutrophils react more strongly in this regard than CD177^neg^ neutrophils could be an indication that CD177 – though not recognized by PR3-ANCA – has, like PR3, a role in facilitating ANCA-initiated neutrophil activation, beyond serving as a platform for additional ANCA epitopes due to complexation with PR3. The experiments presented here address this possibility directly by blocking CD177:PR3 complex formation *in situ* and probing the effect of this epitope removal on the ANCA induced activation response. The clear result was that once the bulk of the CD177 bound PR3 was titrated away from the neutrophil surface, the ANCA sensitivity of the CD177^pos^ population was fully reduced to that of the CD177^neg^ population. ANCA addition still elicited a respiratory burst from these neutrophils, but this via the remaining directly membrane-bound mPR3 that exists in both populations. These results – along with the fact that CD177^neg^ neutrophils are also sensitive to ANCA - make clear that CD177 is not required for the ANCA induced activation effect. They also support the simplest explanation for the enhanced ANCA sensitivity of CD177^pos^ neutrophils, namely that the presence of CD177 – likely all of which is bound to PR3 – makes a larger abundance of ANCA epitopes available than is found on CD177^neg^ neutrophils. CD177 likely ‘presents’ PR3 to the extracellular environment in a uniform orientation that maximizes accessibility to the most common PR3-ANCA epitopes (7). Our results imply that this increase in binding sites is sufficient to account for the increased ANCA sensitivity of these neutrophils, since physically removing them reduces ANCA sensitivity to that of CD177^neg^ neutrophils.

The results do not answer the question of what function(s) CD177 actually has in neutrophils and why it is found in complex with PR3. They also do not rule out entirely that CD177 can participate in the ANCA induced activation process; the fact that multivalent CD177 binders provoke a response similar to that of PR3-ANCA provides clues for further studies into the role of CD177 in neutrophil biology. The newly developed antibodies described here will be of great value in analyzing the function of CD177 in such experiments and may also be of value in the treatment of AAV. Since the removal of CD177-bound PR3 results in a substantial reduction in ANCA-induced neutrophil activation, blocking this interaction *in vivo* could prove beneficial for PR3-AAV patients, particularly those with large CD177^pos^ neutrophil populations.

### Experimental Procedures

#### Hybridoma generation

Recombinant CD177 was prepared as described previously (7) and provided to Biogenes GmbH (Berlin) for inoculation of mice. Hybridomas delivered by Biogenes were cocultured on feeder cells obtained from peritoneal lavage of Black6 mice in hybridoma medium: Dulbecco’s Minimal Essential Medium (DMEM) (Sigma, D5871) supplemented with 20% fetal calf serum (Merck), 2 mM glutamine (Gibco), 1 mM sodium pyruvate (Sigma, S8636) and antibiotics (Pen/Strep, Gibco).

#### IgG isolation and Fab preparation

Stable hybridoma cultures were grown in 300 cm^2^ culture flasks (TPP) in 50 ml hybridoma medium. Cells were split 1:10 every 48 hours (at 80-90% confluence) and with complete exchange into fresh medium. Collected medium aliquots containing IgG were pooled and passed over a 5 ml Protein G Agarose column (GE Healthcare) washed with phosphate buffered saline (PBS, Merck) and eluted with 50 mM glycine (Sigma), 150 mM NaCl (Sigma), pH 3.5 directly into 1 M Tris, pH 8.0 for neutralization. Elution fractions were pooled and concentrated in 100 kD molecular weight cutoff Amicon spin concentrators (Millipore). Concentrated IgG were further purified over a Superdex 200 size exclusion column (GE Healthcare) in 20 mM HEPES (Sigma), 150 mM NaCl, pH 7.4. IgG containing fractions were verified by SDS-PAGE, pooled, concentrated in 30 kD MWCO Amicons and stored at 4°C until use.

Fabs were prepared from purified IgG by incubation with Papain-Agarose (Thermo Fisher) according to the manufacturer’s instructions. Digested IgG was passed over Protein G agarose to remove Fc and undigested IgG and Fab containing fractions were pooled, concentrated in 10 kD MWCO Amicons and subjected to size exclusion chromatography as for the IgG.

#### SPR experiments

Experiments were performed on a ProteOn XPR36 instrument (BioRad) using standard amine chemistry for coupling IgG to the sensor chip (GLH sensor chips, BioRad). Ligand dilution series were prepared in ProteOn running buffer (PBS supplemented with 0.005% Tween-20 (Sigma).

#### Neutrophil preparation

Blood neutrophils from healthy human donors were obtained (Ethic Votum EA1/277/11) and purified as described previously (20). Briefly, neutrophils from healthy volunteers were isolated from heparinized whole blood by red blood cell sedimentation with 1% dextran, followed by Histopaque 1.083 (Sigma) density gradient centrifugation, and hypotonic erythrocyte lysis. Neutrophils were centrifuged and resuspended in HBSS with calcium and magnesium (HBSS++, Merck). Cell viability was determined by Trypan blue exclusion and exceeded 99%.

#### Separation of CD177^pos^/mPR3^high^ and CD177^neg^/mPR3^low^ Neutrophil Subsets by Magnetic Beads

Neutrophil subsets were separated with MACS separation columns (Miltenyi Biotec). Isolated neutrophils were stained with monoclonal anti-PR3 (clone 43-8-3-1). MACS rat anti-mouse IgG1 beads were added, and cells were pipetted onto a MACS LD column and the flowthrough containing the non-labeled CD177^neg^/mPR3^low^ neutrophils was collected. Collumns were removed from the magnet to allow collection of the labeled CD177^pos^/mPR3^high^ cells. The purity of the two separated subsets was assessed by flow cytometry using a FITC-labeled anti-CD177 IgG.

#### Membrane PR3 Expression on Neutrophils

Neutrophils were stimulated with 2 ng/ml TNFα (30 min., 37 °C, R&D systems) to increase the amount of membrane PR3. Cells were washed and stained with monoclonal anti-PR3 (clone 81-3-3)-Alexa488 conjugated IgG (2,5 µg/ml, 20 min. on ice). mPR3 expression was assessed by flow cytometry using a FACSCalibur istrument. Ten thousand events per sample were assayed. For blocking experiments, TNFα -primed neutrophils were incubated with 20 µg/ml anti-CD177 IgG or Fab (60 min., on ice), The capacity to block anti-PR3 IgG binding was tested by subsequent incubation with the Alexa488-conjugated anti-PR3 IgG.

#### Measurement of Respiratory Burst

Superoxide was measured using the assay of SOD-inhibitable reduction of ferricytochrome c. Neutrophils were pretreated with 5 ug/ml cytochalasin B for 15 min. on ice. Cells (0.75 × 10^6^/ ml) were primed with 2 ng/ml TNFα for 15 min. before stimulating antibodies (or Fabs) were added. The final concentration was 5 µg/ml for monoclonal antibodies or Fabs.

For superoxide blocking experiments, cells were incubated with 20 µg/ml or with indicated amounts of anti-CD177 Fab during the TNFα priming, before the stimulating monoclonal anti-PR3 (clone 43-8-3-1) or 75 µg/ml purified PR3-ANCA preparations were added.

Experiments were performed in 96-well plates at 37 °C for up to 45 min., and the absorption of samples with and without 300 U/ml SOD was measured at 550 nm using a microplate reader (Molecular Devices)

## Acknowledgements

This work was funded by Else-Kröner-Fresenius Stiftung Grant 2013_A73 to R.K. and S.F.M. and by the Deutsche Forschungsgemeinschaft (project number 246135856) to R.K.

## References

1. Borregaard, N. (2010) Neutrophils, from marrow to microbes. Immunity 33, 657–670

2. Vogt, K. L., Summers, C., Chilvers, E. R., and Condliffe, A. M. (2018) Priming and depriming of neutrophil responses in vitro and in vivo. Eur J Clin Invest, e12967

3. Campbell, E. J., Campbell, M. A., and Owen, C. A. (2000) Bioactive proteinase 3 on the cell surface of human neutrophils: quantification, catalytic activity, and susceptibility to inhibition. J Immunol 165, 3366–3374

4. Halbwachs-Mecarelli, L., Bessou, G., Lesavre, P., Lopez, S., and Witko-Sarsat, V. (1995) Bimodal distribution of proteinase 3 (PR3) surface expression reflects a constitutive heterogeneity in the polymorphonuclear neutrophil pool. FEBS Lett 374, 29–33

5. von Vietinghoff, S., Tunnemann, G., Eulenberg, C., Wellner, M., Cristina Cardoso, M., Luft, F. C., and Kettritz, R. (2007) NB1 mediates surface expression of the ANCA antigen proteinase 3 on human neutrophils. Blood 109, 4487–4493

6. Bauer, S., Abdgawad, M., Gunnarsson, L., Segelmark, M., Tapper, H., and Hellmark, T. (2007) Proteinase 3 and CD177 are expressed on the plasma membrane of the same subset of neutrophils. J Leukoc Biol 81, 458–464

7. Jerke, U., Marino, S. F., Daumke, O., and Kettritz, R. (2017) Characterization of the CD177 interaction with the ANCA antigen proteinase 3. Sci Rep 7, 43328

8. Witko-Sarsat, V., Lesavre, P., Lopez, S., Bessou, G., Hieblot, C., Prum, B., Noel, L. H., Guillevin, L., Ravaud, P., Sermet-Gaudelus, I., Timsit, J., Grunfeld, J. P., and Halbwachs-Mecarelli, L. (1999) A large subset of neutrophils expressing membrane proteinase 3 is a risk factor for vasculitis and rheumatoid arthritis. J Am Soc Nephrol 10, 1224–1233

9. Schreiber, A., Busjahn, A., Luft, F. C., and Kettritz, R. (2003) Membrane Expression of Proteinase 3 Is Genetically Determined. J Am Soc Nephrol 14, 68–75

10. Eulenberg-Gustavus, C., Bahring, S., Maass, P. G., Luft, F. C., and Kettritz, R. (2017) Gene silencing and a novel monoallelic expression pattern in distinct CD177 neutrophil subsets. J Exp Med 214, 2089–2101

11. Rarok, A. A., Stegeman, C. A., Limburg, P. C., and Kallenberg, C. G. M. (2002) Neutrophil Membrane Expression of Proteinase 3 (PR3) Is Related to Relapse in PR3-ANCA-Associated Vasculitis. J Am Soc Nephrol 13, 2232–2238

12. Schreiber, A., Otto, B., Ju, X., Zenke, M., Goebel, U., Luft, F. C., and Kettritz, R. (2005) Membrane proteinase 3 expression in patients with Wegener’s granulomatosis and in human hematopoietic stem cell-derived neutrophils. J Am Soc Nephrol 16, 2216–2224

13. Hu, N., Westra, J., Huitema, M. G., Bijl, M., Brouwer, E., Stegeman, C. A., Heeringa, P., Limburg, P. C., and Kallenberg, C. G. (2009) Coexpression of CD177 and membrane proteinase 3 on neutrophils in antineutrophil cytoplasmic autoantibody-associated systemic vasculitis: anti-proteinase 3-mediated neutrophil activation is independent of the role of CD177-expressing neutrophils. Arthritis and rheumatism 60, 1548–1557

14. Abdgawad, M., Gunnarsson, L., Bengtsson, A. A., Geborek, P., Nilsson, L., Segelmark, M., and Hellmark, T. (2010) Elevated neutrophil membrane expression of proteinase 3 is dependent upon CD177 expression. Clin Exp Immunol 161, 89–97

15. Schreiber, A., Luft, F. C., and Kettritz, R. (2004) Membrane proteinase 3 expression and ANCA-induced neutrophil activation. Kidney Int 65, 2172–2183

16. Kettritz, R., Jennette, J. C., and Falk, R. J. (1997) Crosslinking of ANCA-antigens stimulates superoxide release by human neutrophils. J Am Soc Nephrol 8, 386–394

17. Kuckleburg, C. J., and Newman, P. J. (2013) Neutrophil proteinase 3 acts on protease-activated receptor-2 to enhance vascular endothelial cell barrier function. Arterioscler Thromb Vasc Biol 33, 275–284

18. van den Berg, C. W., Tambourgi, D. V., Clark, H. W., Hoong, S. J., Spiller, O. B., and McGreal, E. P. (2014) Mechanism of neutrophil dysfunction: neutrophil serine proteases cleave and inactivate the C5a receptor. J Immunol 192, 1787–1795

19. Padrines, M., Wolf, M., Walz, A., and Baggiolini, M. (1994) Interleukin-8 processing by neutrophil elastase, cathepsin G and proteinase-3. FEBS Lett 352, 231–235

20. Schreiber, A., Pham, C. T., Hu, Y., Schneider, W., Luft, F. C., and Kettritz, R. (2012) Neutrophil Serine Proteases Promote IL-1beta Generation and Injury in Necrotizing Crescentic Glomerulonephritis. Journal of the American Society of Nephrology : JASN 23, 470–482

21. Witko-Sarsat, V., Canteloup, S., Durant, S., Desdouets, C., Chabernaud, R., Lemarchand, P., and Descamps-Latscha, B. (2002) Cleavage of p21waf1 by proteinase-3, a myeloid-specific serine protease, potentiates cell proliferation. The Journal of biological chemistry 277, 47338–47347

22. Jerke, U., Hernandez, D. P., Beaudette, P., Korkmaz, B., Dittmar, G., and Kettritz, R. (2015) Neutrophil serine proteases exert proteolytic activity on endothelial cells. Kidney Int 88, 764–775

23. Vong, L., D’Acquisto, F., Pederzoli-Ribeil, M., Lavagno, L., Flower, R. J., Witko-Sarsat, V., and Perretti, M. (2007) Annexin 1 cleavage in activated neutrophils: a pivotal role for proteinase 3. The Journal of biological chemistry 282, 29998–30004

24. Kantari, C., Millet, A., Gabillet, J., Hajjar, E., Broemstrup, T., Pluta, P., Reuter, N., and Witko-Sarsat, V. (2011) Molecular analysis of the membrane insertion domain of proteinase 3, the Wegener’s autoantigen, in RBL cells: implication for its pathogenic activity. J Leukoc Biol 90, 941–950

